# Effect of ligands on HP-induced unfolding and oligomerization of β-lactoglobulin

**DOI:** 10.1101/2020.06.29.177972

**Authors:** S. Minić, B. Annighöfer, A. Hélary, D. Hamdane, G. Hui Bon Hoa, C. Loupiac, A. Brûlet, S. Combet

## Abstract

To probe intermediate states during unfolding and oligomerization of proteins remains a major challenge. High pressure (HP) is a powerful tool for studying these problems, revealing subtle structural changes in proteins not accessible by other means of denaturation. Bovine β-lactoglobulin (BLG), the main whey protein, has a strong propensity to bind various bioactive molecules, such as retinol and resveratrol, two ligands with different affinity and binding sites. By combining *in situ* HP-small-angle neutron scattering (SANS) and HP-UV/visible absorption spectroscopy, we report the specific effects of these ligands on 3D conformational and local changes in BLG induced by HP. Depending on BLG concentration, two different unfolding mechanisms are observed *in situ* under pressures up to ~300 MPa, mediated by the formation of disulfide bonds: either a complete protein unfolding, from native dimers to Gaussian chains, or a partial unfolding with oligomerization in tetramers. Retinol, which has a high affinity for BLG hydrophobic cavity, significantly stabilizes BLG both in 3D and local environments, by shifting the onset of protein unfolding by ~100 MPa. Increasing temperature from 30 to 37°C enhances the hydrophobic stabilization effects of retinol. In contrast, resveratrol, which has a low binding affinity for site(s) on the surface of the BLG, does not induce any significant effect on the structural changes of BLG due to pressure. HP treatment back and forth up to ~300 MPa causes irreversible covalent oligomerization of BLG. *Ab initio* modeling of SANS shows that the oligomers formed from BLG/retinol complex are smaller and more elongated compared to BLG without ligand or in the presence of resveratrol. By combining HP-SANS and HP-UV/vis absorption spectroscopy, our strategy highlights the crucial role of BLG hydrophobic cavity and opens up new possibilities for the structural determination of HP-induced protein folding intermediates and irreversible oligomerization.

**STATEMENT OF SIGNIFICANCE:** High pressure (HP) is a powerful probe to access the intermediate states of proteins through subtle structural changes not accessible by other means of denaturation. Bovine β-lactoglobulin (BLG), the main whey protein, is able to bind various bioactive molecules, such as retinol and resveratrol, exhibiting different affinity and binding sites. By combining HP-small-angle neutron scattering and HP-UV/visible absorption spectroscopy, we highlight two different mechanisms during the unfolding and oligomerization of BLG depending on protein concentration. Above all, we show that retinol significantly prevents the unfolding and oligomerization of BLG, unlike resveratrol, emphasizing the crucial role of the hydrophobic cavity in BLG stabilization. Our strategy opens up new possibilities for the structural determination of protein intermediates and oligomers using HP.

## INTRODUCTION

The understanding of protein folding, misfolding, and unfolding remains a major challenge in structural biology. High pressure (HP) is a powerful tool to probe the mechanisms of protein folding by elucidating the dynamics and structure of folding intermediates (1). Nowadays, it is very well-known that the use of HP results in the disruption of the native structure of proteins due to the decrease of the volume of the protein/solvent complex upon denaturation. HP studies thus provide a fundamental thermodynamic parameter for protein unfolding, *i.e*. the volume change (2). Recent studies demonstrate that pressure mostly unfolds proteins through hydrophobic cavities present in the folded state that are eliminated in the unfolded states (3). Internal cavities are thus important structural features for proteins and sources of fluctuation between different conformational states, which can be stabilized by subsequent ligand binding.

Bovine β-lactoglobulin (BLG), the main whey protein of cow milk, belongs to the lipocalin protein family, whose members folds up into an eight stranded antiparallel β-barrels, arranged to form the central hydrophobic cavity (4) (Fig. 1). An intramolecular disulfide bridge between cysteins (Cys) 106 and 119 stabilizes this β-barrel structure. Additionally, Cys66 and Cys160 residues form the second disulfide bond, close to the edge of the hydrophobic cavity (5), while Cys121 exists in free form and is buried under an α-helix located on the surface of the native protein (6) (Fig. 1). The presence of a hydrophobic cavity gives BLG the ability to bind reversibly with high affinity various hydrophobic ligands such as retinol (vitamin A, owning mostly a hydrophobic tail), other fatty acids, cholesterol, vitamin D, etc. (4). In contrast, other ligands, such as bioactive polyphenol resveratrol (7) and protoporphyrin IX (8), much less hydrophobic, could bind surface site(s) of BLG but with a lower affinity. In the present study, we focus on retinol and resveratrol, these two BLG ligands exhibiting (*i*) different BLG binding sites (Fig. 1), (*ii*) high or low affinity, with *K_a_* of 10^8^ and 10^4^ L/mol for retinol and resveratrol, respectively (4,7), and (*iii*) a bit higher solubility at pH 7 for resveratrol than for retinol.

**Figure 1.**
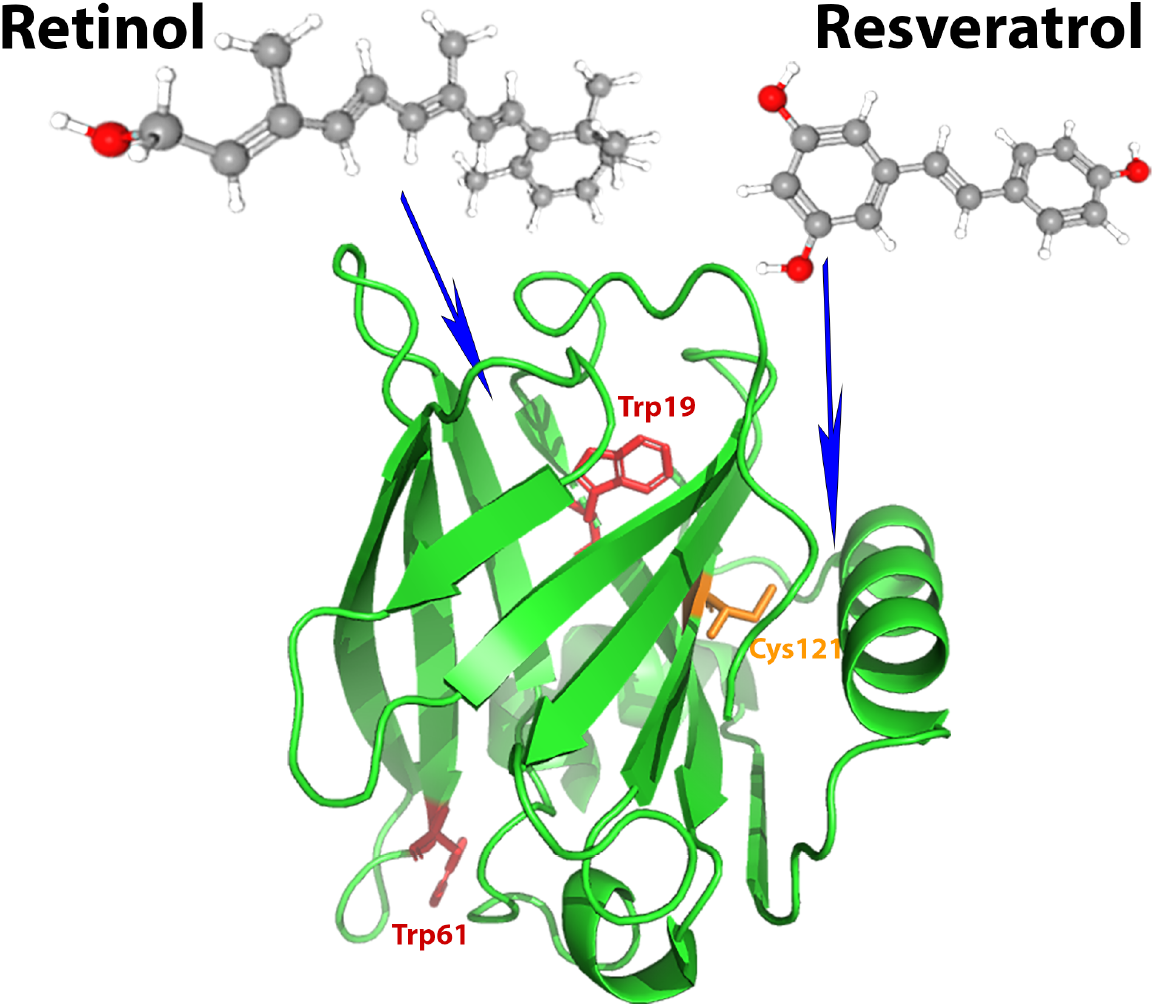
Ribbon model of the crystal structure of BLG (1GX8.pdb). Arrows indicate retinol (18) and resveratrol (19) approximative binding sites. Tryptophan (Trp) aromatic side chains are shown in red (Trp19 and Trp61), with Trp19 residue close to the BLG cavity. The free cystein Cys121, colored in orange, is non accessible in the native folded state of BLG but is exposed to the solvent after protein unfolding.

BLG behavior at HP conditions has been extensively studied by numerous techniques, such as UV/visible (UV/vis) absorption (9), fluorescence (10), and Fourier-transform infra-red (FT-IR) spectroscopy (11), as well as 2D nuclear magnetic resonance (NMR) (12) and small-angle X-ray or neutron scattering (SAXS or SANS, respectively) (11) (13). BLG exists in the native state as monomer at pH 2 and dimer at neutral pH (14). Due to its hydrophobic cavity, BLG is very sensitive to pressure denaturation and thereby constitutes an adequate model to study HP protein unfolding. It unfolds at pressures between 100-200 MPa (11) but pressure stability of the protein strongly depends on its solvent pH, the protein being more stable at pH 2 than at neutral pH (10). It is well-known that ligand binding can prevent the thermal denaturation of proteins (15). However, the effect of ligands on the pressure-dependent structural changes of proteins have not been thoroughly studied, compared to thermal or chemical denaturation. From a fundamental point of view, BLG represents therefore a relevant and sensitive model system to study the effect of ligand binding on protein stability, folding, and self-association (16) under HP conditions. BLG is able to form non-covalent oligomers and such an oligomerization strongly depends on pH and ionic strength (14). An increased population of unfolded BLG molecules can also promote amyloid fibril formation (17).

In the present study, we compare the effects of retinol and resveratrol ligands on HP-dependent stability of BLG. We measured both *in situ* HP-SANS, to study the tridimensional conformational changes, and *in situ* HP-UV/vis absorption, to follow the changes in the local environment of aromatic amino acid residues (tryptophan (Trp) residues, especially) (Fig. 1). We report that retinol strongly reduces HP-induced large conformational changes on BLG, whereas resveratrol binding does not produce any significant changes. HP treatment back and forth up to ~300 MPa causes irreversible BLG covalent oligomerization, while keeping “molten globule” folded conformations. The high affinity binding of retinol in BLG hydrophobic cavity prevents such protein oligomerization, in contrast to resveratrol.

## MATERIALS & METHODS

### Materials

BLG was purified as previously described by Fox *et al*. (20). Protein concentration was determined spectrophotometrically using the extinction coefficient of 17,600 M^−1^ cm^−1^ at 278 nm (21). BLG was dialyzed against 50 mM Tris buffer in D_2_O (pD 7.2) or 100 mM Tris buffer in H_2_O (pH 7.2) for HP-SANS and HP-UV/vis measurements, respectively.

Ligands were purchased from Sigma Aldrich (USA). Retinol and resveratrol were solubilized in deuterated ethanol (d6, Sigma Aldrich) with a concentration not exceeding 2% (*v*/*v*). The stoechiometry of both ligands is stated to be 1:1 protein:ligand molar ratio (19). In SANS measurements, due to very low solubility, the hydrophobic retinol ligand was used at only 5 mg/mL, whereas resveratrol was solubilized at 8 mg/mL. All measurements were performed at 30°C, unless otherwise stated, and at pH/pD 7.2 to enhance ligand solubility, especially for retinol. Fully chemical unfolding of BLG was performed by dissolving the protein in D_2_O solution of 6 M deuterated guanidine in DCl (Sigma Aldrich) at pD adjusted to 7.2. All other chemicals were of analytical reagent grade and milli-*Q* water was used as solvent.

### SDS-PAGE

Sodium dodecyl sulfate-polyacrylamide gel electrophoresis of BLG, in the absence or presence of ligands, before and after HP-SANS measurements, was performed under nonreducing conditions (22), unless it is otherwise stated. An amount of 12 μg of each protein sample was applied on 4-20% gradient precast gel (BioRad, USA). Gels were stained using Coomassie brilliant blue G-250 (Sigma Aldrich) and band intensities were quantitated using ImageJ software (National Institutes of Health, USA). The intensity of each band was normalized to the total band intensity within sample and results were expressed as %.

### HP-UV/visible absorption spectroscopy

UV/vis absorption spectra under *in situ* HP were recorded on a Cary 3E spectrometer (Varian, USA) using HP optical bomb with sapphire windows and HP generator as previously described (23). Square quartz cell (with an optical pathlength of 5 mm) containing the sample was positioned within HP optical bomb, while a plastic membrane on the top of the cell separates the sample from pressure transmitting liquid (H_2_O). Absorbance of BLG solution (100 μM or 2 mg/mL) in the presence or the absence of 100 μM ligand (retinol or resveratrol) was recorded between 250 and 310 nm, with a bandwidth of 1 nm and a data interval of 0.1 nm, at a scanning speed of 30 nm/min. Spectra were recorded at various pressures (between 0.1 to 300 MPa, with steps of 20 or 30 MPa), at 30 or 37°C. The pressure was increased at a speed of 10 MPa/min. Measurements of buffer solutions in the absence or presence of ligands (without protein) were performed at the same above conditions and recorded spectra were subtracted from the spectra of BLG without ligand or BLG/ligand complexes, respectively. The fourth derivative spectra were calculated by OriginPro 8.5 Software (USA) using Savitzky-Golay smoothing algorithm with 100-point window. The exact position of the maximum absorption bands at ~291 nm (characteristic of Trp residues of BLG) in the fourth derivative spectra was determined and the percentage of protein unfolding was determined using the following equation:

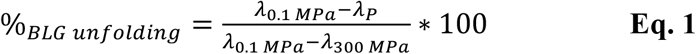

where *λ*_0.1 MPa_, *λ*_300 MPa_, and *λ_P_* represent the absorption maximum at 0.1 MPa, 300 MPa, and at a given pressure, respectively. Absorption at 300 MPa is used as a reference of unfolded BLG protein, as shown by Dufour *et al*. (10). The results were expressed as the percentage of unfolded BLG as a function of pressure. The Gibbs free energy (Δ*G*_0.1 *MPa*_ at 0.1 MPa) and the apparent volume change of unfolding (Δ*V_u_*) were determined by fitting the pressure denaturation curves with the following equation adapted from (24), by replacing the *A_l_* and *A_h_* amplitudes by 0 and 100, respectively:

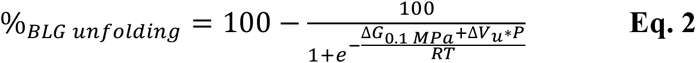

The pressure value at which one half of proteins is unfolded (50%) represents the half denaturation pressure (*P_1/2_*) of BLG.

### HP-SANS measurements

SANS measurements were performed on PACE SANS instrument at the Laboratoire Léon-Brillouin (LLB, Saclay, France). HP-SANS experiments were performed using a new HP-SANS cell developed at LLB (25), using two metallic TiAl6V4 ELI windows (3 mm-thickness each) and a sample pathlength of 4.6 mm. HP-SANS spectra were recorded at 5, 100, 150, 200, 250, and 300 MPa. The scattering signals of empty HP cell, empty beam, and electronic noise were also recorded to be able to extract the scattering signal from BLG solutions despite the large scattering signal coming from empty HP cell (26) (Fig. S1). Each BLG sample was measured in a quartz Hellma^®^ cell (with a path length of 2 mm) before and after HP treatment. For measurements in quartz cell, the *Q*-range covers 6.0 10^−3^ - 0.5 Å^−1^, whereas for measurements in HP cell, due to geometrical and scattering constraints, the *Q*-range was limited to 2 10^−2^ - 0.2 Å^−1^. However, this latter *Q*-range is adequate to analyse BLG structural changes.

BLG/retinol and BLG/resveratrol complexes (1:1 molar ratio) were measured at protein dilute concentrations to limit aggregation, at 5 mg/mL (274 μM) or 8 mg/mL (440 μM), respectively. The reduced protein concentration of retinol/BLG complex, due to the lower solubility of retinol compared to resveratrol, was used to avoid ligand precipitation. BLG protein without ligand was measured at the same concentration, as a control, containing 2% (*v/v*) deuterated ethanol. Pressure was increased at a rate of about 10 MPa/min. Measurements were performed at 30 or 37°C for retinol to compare with a condition with both higher ligand solubility and binding affinity. The pressure-dependence for buffer SANS intensity was also checked to be negligible (not shown). A constant background of 0.05-0.06 cm^−1^, deduced by measuring scattering intensity in a Hellma^®^ cell at the *Q*-range between 0.4 and 0.5 Å^−1^, was subtracted, as incoherent signal, from the SANS intensities of all BLG solutions under pressure.

Air bubbles, that may be introduced during the sample injection into the HP cell, were compressed by increasing gently the pressure up to 5 MPa. We checked that BLG SANS curves in the HP cell at this “ambient” pressure were mostly superimposable to the curves of the same sample measured in a Hellma^®^ cell (Fig S1), showing that SANS curves at 5 MPa can be considered as references for native BLG.

### SANS data analysis

The classical expression of the scattering intensity *I*(*Q*) (in cm^−1^) of spherically symmetric, homogenous, and relatively monodisperse particles can be written:

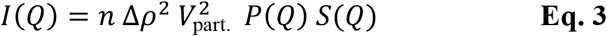

where *n* is the number of particles *per* volume unit (cm^−3^), Δ*ρ* the difference of the neutron scattering length density between the particles and the solvent (cm^−2^), and *V*_part._ (cm^3^) the specific volume of the particles.

The form factor *P*(*Q*) describes the shape of the particles and fulfills the condition *P*(0) = 1, while the structure factor *S*(*Q*) describes the interactions between the particles. In the absence of interactions, like in a dilute solution, *S*(*Q*) = 1. So, from the scattered intensity *I*(0), we can extract the mass of the particle in atomic units (*MW*, in g/mol), which coincides with the molecular mass if the particle consists of a single molecule, by introducing the concentration 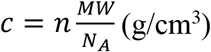 and the particle density 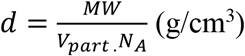 using the following equation:

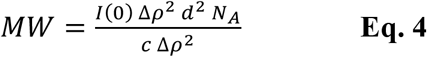

where *N_A_* is the Avogadro number and *Δρ*^2^ the contrast.

ScÅtter software (http://www.bioisis.net/tutorial/9) was used to determine the intensity at zero angle *I*(0) and the radius of gyration (*R*g). These values are defined at small *Q*-values (*QRg* < 0.8-1.4) by the Guinier approximation with (27):

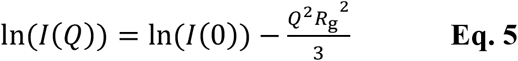

The distance distribution function *P*(r) was calculated with PRIMUS (ATSAS) program (28).

*In situ* HP-SANS data were fitted using a cylinder model (27) in SasView software (https://www.sasview.org), with obtained chi^2^ values not exceeding 1.2. However, the SANS data measured after HP treatment for BLG oligomers were better fitted with a triaxial ellipsoid model (29) with chi^2^ values given in Table 3. BLG oligomers were also modeled by *ab initio* rigid body modeling using MASSHA software (ATSAS) (30): oligomers were built using crystal structures from, either BLG monomer (3BLG.pdb), or BLG dimer (1BEB.pdb), as starting building blocks. The previously best fitted ellipsoid dimensions were taken into account to estimate the number and position of monomers or dimers, which were further adjusted until a satisfactory fit (chi^2^ values given in Table 3).

## RESULTS

### Structural features for native and fully unfolded BLG

We checked that native BLG at pD 7.2 is dimeric by fitting SANS data with the theoretical curve obtained from the dimeric BLG crystal structure (1GX8.pdb (18)) using CRYSON software (ATSAS) (31) (Fig. 2). Besides, using the scattering length density of BLG determined by “contrast variation” experiment (Table 1), we estimated, using Eq. 4 with *d* = 1.35 g/cm^3^ and *c* = 8 mg/mL, that *MW* of the scattering particles is equal to 37,924 ± 1,129 g/mol, in very good agreement with the theoretical *MW* of dimeric BLG (37,142 g/mol). The pair-distribution function *P*(r) analysis, giving a *D*_max_ of ~71 Å, confirms the dimeric conformation of BLG in our study conditions (Fig. 2B).

**Figure 2.**
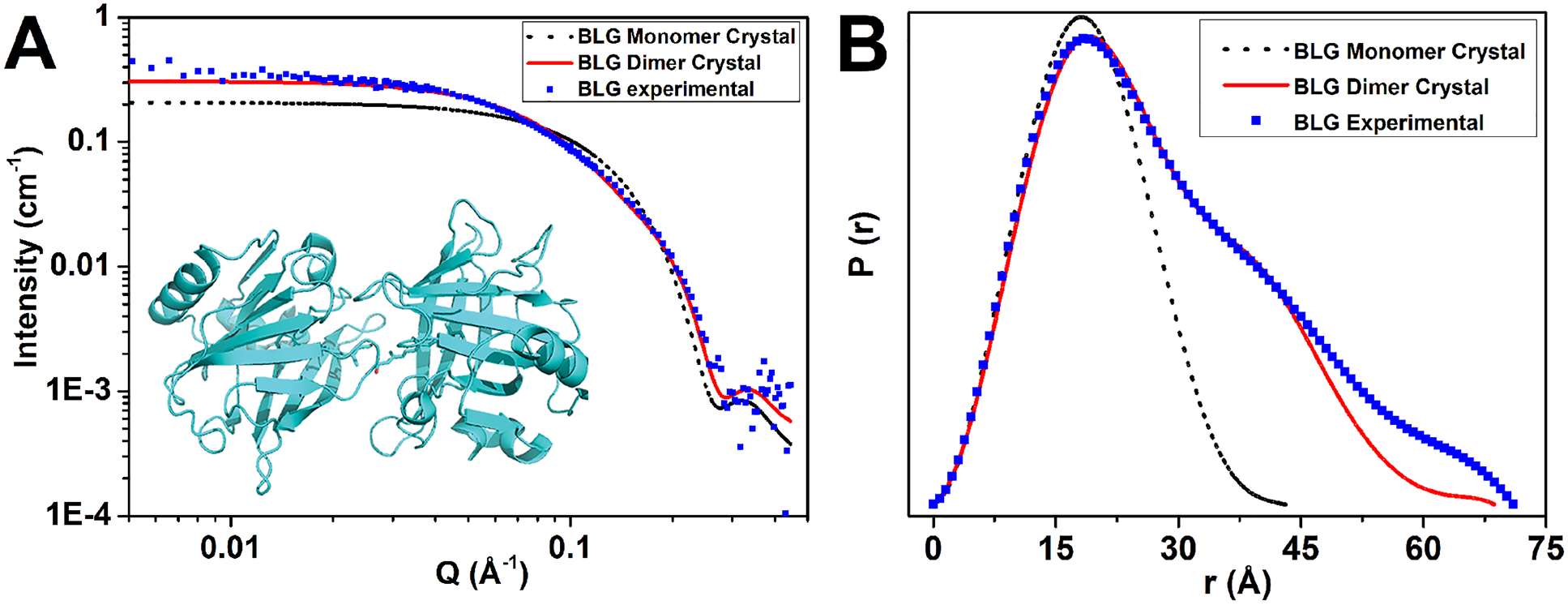
(***A***) SANS intensities of BLG (8 mg/mL) in 50 mM Tris D_2_O buffer at pD 7.2. The full red and dotted black lines correspond to the theoretical curves calculated from BLG dimer or monomer crystal structures, respectively, using CRYSON software with 1GX8.pdb as monomer or dimer crystallographic file. Insert is a representation of BLG dimer (1GX8.pdb). (***B***) The pair-distribution function *P*(r) analysis with the same curve code, showing *D*_max_ of ~71 Å for BLG.

**Table 1.** Molecular characteristics, scattering length, and scattering length density of BLG.

| | Molecular weight<br>($MW$, g/mol) | Density<br>($d$, g/cm<sup>3</sup>) | Scattering<br>length<br>(10<sup>-12</sup> cm) | Scattering length density<br>($\rho$, 10<sup>10</sup> cm<sup>-2</sup>) | |
| --- | --- | --- | --- | --- | --- |
| | | | | $\rho_{theo}$ | $\rho_{exp}^*$ |
| BLG | 37,142 (dimer) | 1.34 | 807 | 1.81 | 2.72 |
| D<sub>2</sub>O | 20.027 | 1.10 | 1.91 | 6.38 | / |

\* $\rho_{exp}$ is deduced from contrast variation measurements on BLG using different H<sub>2</sub>O/D<sub>2</sub>O mixtures for protein buffer, giving a matching point at $\Phi(\text{D}_2\text{O}) = 38.5\%$ v/v, thus a scattering length density $\rho_{exp}$ of 2.72 10<sup>10</sup> cm<sup>-2</sup>. The percentage of H-D exchange is deduced to be equal to 79.7%. The theoretical value, $\rho_{theo}$ , is calculated without hydrogen exchange.

The presence of deuterated 6 M guanidine (Gdn)-DCl completely unfolds BLG, as shown by the SANS data that fit the Gaussian coil model commonly used for fully unfolded proteins (Fig. S2) (32). The radius of gyration (*R_g_*) increases from 24 ± 1 Å for native BLG to 41 ± 4 Å for the fully chemically unfolded protein. The bell-shaped plot in Kratky representation confirms the well-folded and compact state of native BLG, while a “plateau” characteristic of Gaussian chains is observed in the presence of Gdn-DCl (Fig. S2B). In native conditions, BLG/retinol and BLG/resveratrol complexes are superimposed in SANS measurements to BLG without ligand (not shown).

### Effect of high pressure on BLG structure

SANS and UV/vis absorption spectroscopy measurements were performed by applying *in situ* HP up to ~300 MPa on BLG without ligand compared to BLG/retinol and BLG/resveratrol complexes.

For BLG without ligand and both BLG/ligand complexes, no significant change in the protein 3D conformation is observed by HP-SANS up to ~150 MPa, whether for retinol (Fig. 3) or for resveratrol (Fig. S3). For such pressures, *R_g_* values (Fig. 4, B & D), as well as the length and radius parameters extracted from the cylinder model used to fit SANS data (Fig. 4, A & C) are similar compared to those obtained at the atmospheric pressure. Values of *I*_*Q*→0_, reflecting protein *MW* and hydration-dependent contrast value (see Eq. 4), decrease very slightly with pressure (Fig. S4).

**Figure 3.**
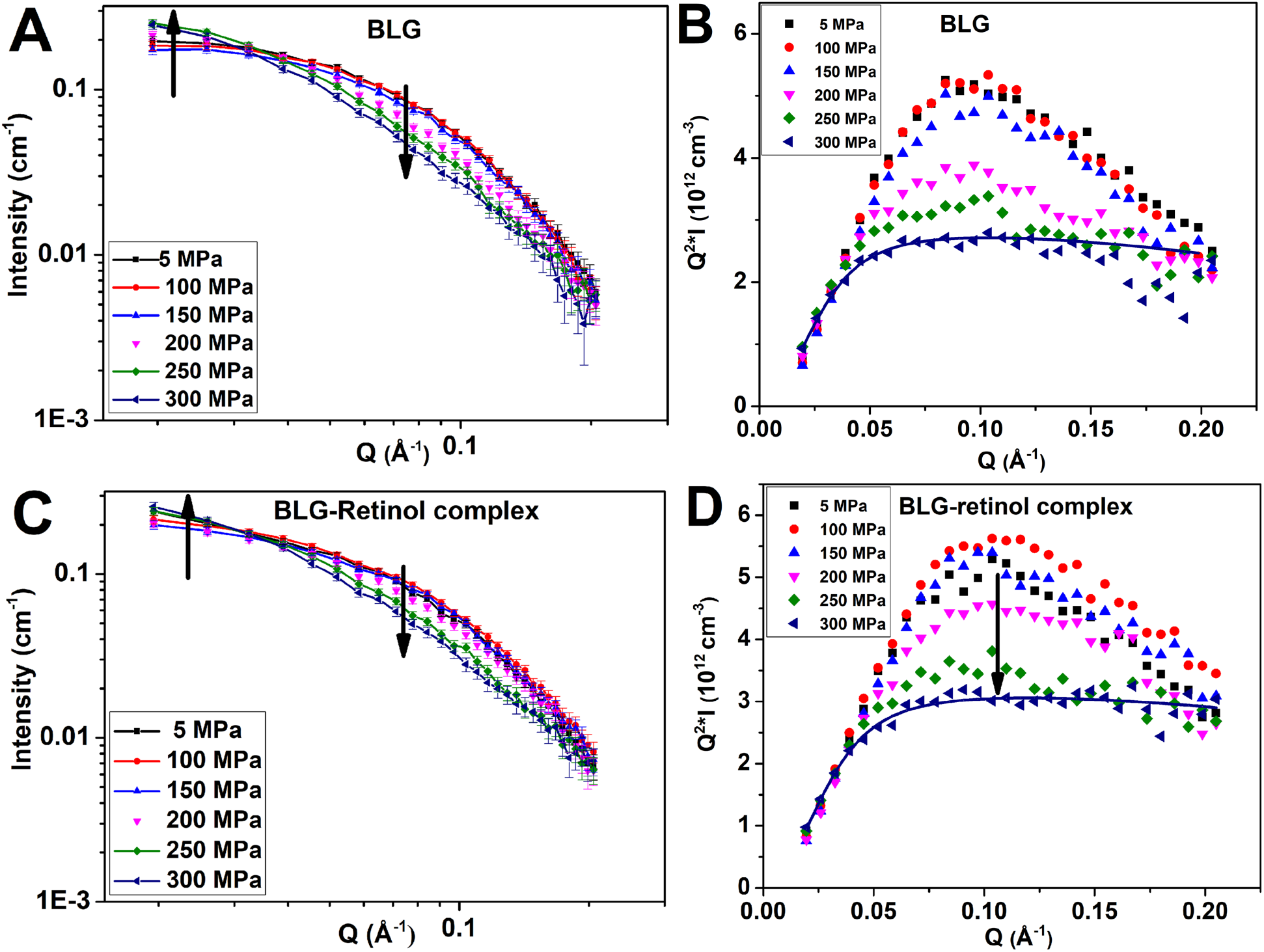
(***A***, ***C***) *In situ* HP-SANS intensities of BLG (5 mg/mL, pD 7.2, 30°C) in the absence or presence of retinol, at different pressures. The arrows represent the directions of SANS intensity changes with increasing pressure. Full lines are guides for the eyes. (***B***, ***D***) The same data in Kratky representation. The dark blue full lines represent the fits obtained at ~300 MPa, with the “plateau” characteristic of a Gaussian chain conformation. For BLG without ligand, fitting has been performed up to ~0.12 Å, due to noisy data at higher *Q*-values.

**Figure 4.**
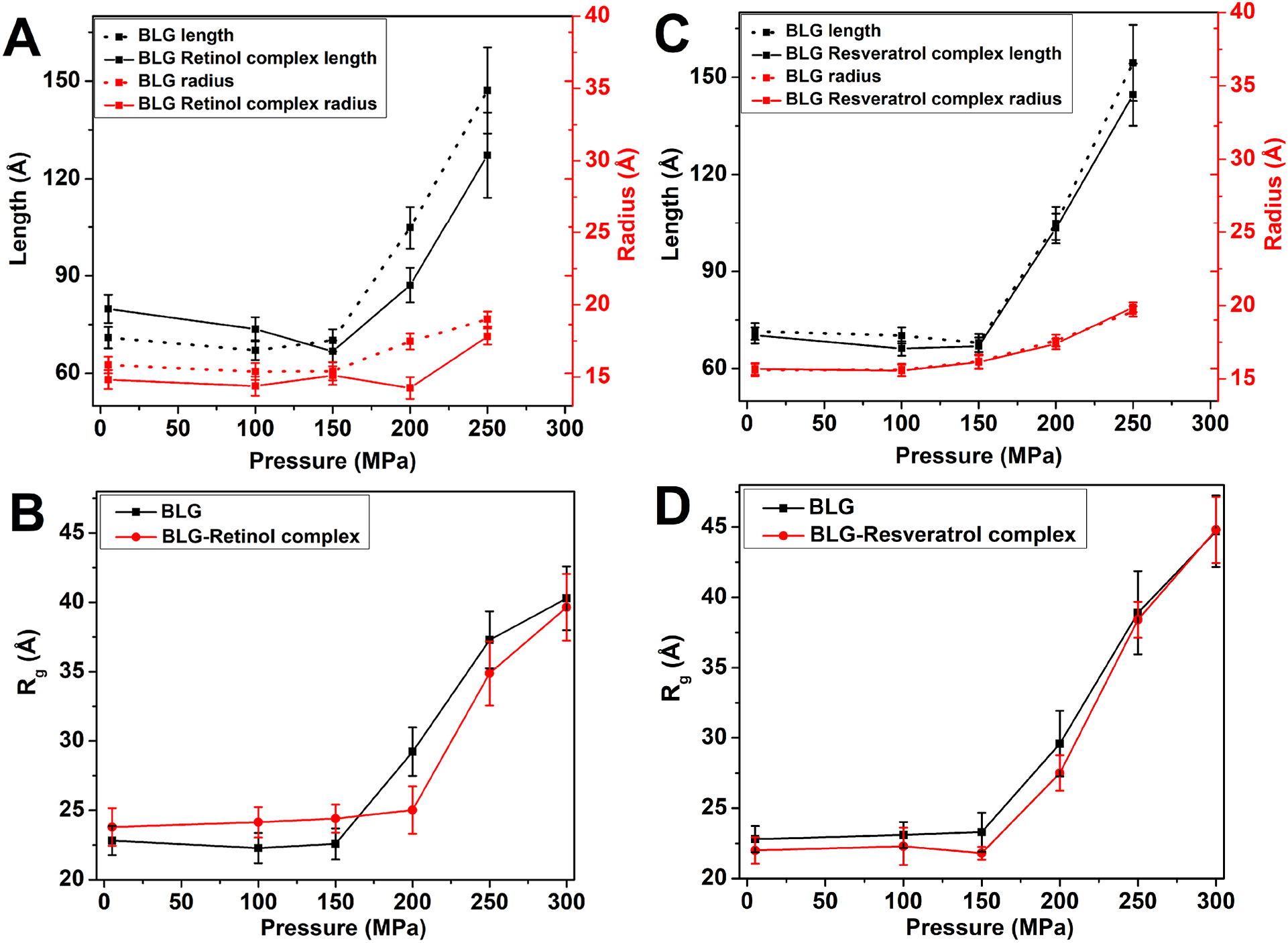
Effects of ligand binding on HP-induced 3D conformational and Trp local unfolding of BLG, at 30°C. (***A***, ***C***) Evolution as a function of pressure of the cylinder length and radius obtained from the analytical model that fits HP-SANS data, in the absence or presence of (*A*) retinol (5 mg/mL BLG) or (*C*) resveratrol (8 mg/mL BLG). At ~300 MPa, the virtually fully unfolded protein cannot be fitted by a cylinder form factor anymore. (***B***, ***D***) Dependence as a function of pressure of the radius of gyration obtained from HP-SANS data, in the absence or presence of (*B*) retinol (5 mg/mL BLG) or (*D*) resveratrol (8 mg/mL BLG).

At higher pressures, two different transitions are observed according to protein concentration. At 5 mg/mL (Fig. 3, A-B), BLG starts to unfold continuously from ~200 MPa to reach a Gaussian chain conformation at ~300 MPa, without any significant change in the *MW* of the dimeric protein. This is clearly observed in Kratky representation with a transition from bell-shaped curves, without any change in the *Q*-position of the peak, to a “plateau”, characteristic of *Q*^−2^ curves without any defined form factor nor volume (Fig. 3B). BLG conformational changes are highlighted by both higher *I*_*Q*→0_ (Fig. S4A) and *R_g_* (Fig. 4B) to values up to ~40 Å, found for the fully chemically unfolded protein. In contrast, at 8 mg/mL, BLG unfolding from ~200 MPa is not complete since the bell-shaped curves in Kratky representation decrease but still tend to zero, and not a constant, at high *Q*-values (Fig. S3B). At that concentration, we observe a transition from a globular state to a larger one, as shown by the shift of the curve peak to lower *Q*-values (Fig. S3B). BLG oligomerizes from the native dimer observed up to ~150 MPa to a tetrameric conformation at ~250-300 MPa, as shown by the doubled *I*_*Q*→0_ value (Fig. S4B), as well as higher *R_g_* (Fig. 4D). For both 5 and 8 mg/mL, the SANS curves are virtually superimposed up to ~150 MPa, as well as at ~250-300 MPa, whereas, in between, the curves at ~200 MPa may illustrate the coexistence of two protein populations (Figs. 3 and S3). SANS curves can be fitted by a cylinder form factor up to ~150 MPa (Fig. 4, A & C). At ~200-250 MPa, this model still fits the data but such a fitting appears trickier, especially at 5 mg/mL, for which BLG tends to a Gaussian polymer chain (Fig. 4, A & C). At ~300 MPa, the cylinder model used as a form factor does not fit the data anymore at both BLG concentrations.

### Effect of ligands on HP-induced structural changes of BLG

Retinol, which binds with high affinity to BLG hydrophobic cavity (18), shifts the onset of the pressure unfolding by ~50 MPa (Fig. 3, C-D). From ~200 MPa, the length and radius of the cylinder form factor, *R_g_*, and *I*_*Q*→0_ are reduced by ~20% compared to BLG without ligand (Figs. 4, A-B, and S4A). Above ~200 MPa, BLG/retinol complex starts to unfold and shows a large increase of the cylinder length and radius parameters, *R_g_*, and *I*_*Q*→0_, approaching the values found for BLG without ligand at ~300 MPa (Figs. 4, A-B, and S4A).

In contrast, resveratrol, with low affinity to BLG surface site(s) (19, 33), does not induce any significant change of the structural features reported above for BLG without ligand from *in situ* HP-SANS (Fig. S3). However, the transition with pressure described for BLG without ligand at 8 mg/mL appears slightly more “abrupt” in the presence of resveratrol, with two well marked protein states, one up to ~150 MPa and the other at ~250-300 MPa, with a transitory state at ~200 MPa (Fig. S3D).

In HP-UV/vis absorption study, using the 4^th^ derivative mode, BLG spectra exhibit absorbance maxima at 276, 291, and 284 nm, corresponding to tyrosine (Tyr) or tryptophan (Trp) residues or both of them, respectively (Figs. 5, A-B, and S5). For all BLG samples, HP treatment induces a “blue shift” of these maxima, especially of the Trp-dependent peak, towards lower wavelengths, indicating local unfolding of the protein (9) (Figs. 5 and S5).

**Figure 5.**
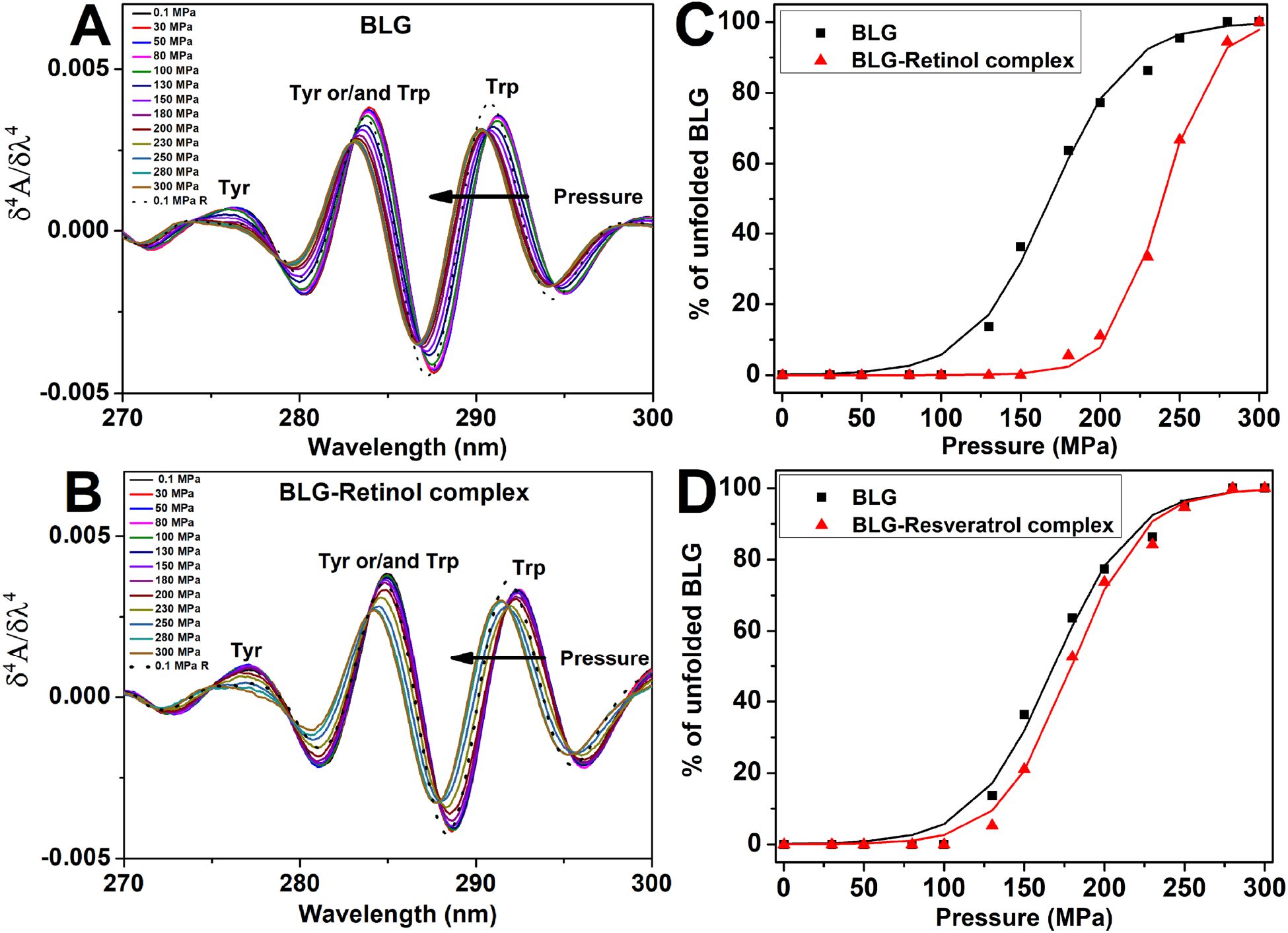
(***A***, ***B***) 4^th^ derivative of *in situ* HP-UV/vis absorption spectra of BLG (2 mg/mL, pH 7.2, 30°C), as a function of pressure, in (*A*) the absence or (*B*) the presence of retinol (1:1 protein:ligand molar ratio). “0.1 MPa R” is the curve obtained by decreasing back the pressure to 0.1 MPa after ~300 MPa treatment. (***C***, ***D***) Percentage of unfolded BLG (Eq. 1) and the corresponding unfolding fits (full lines, Eq. 2), as a function of pressure, in the absence or presence of (*C*) retinol or (*D*) resveratrol, obtained from the 4^th^ derivative mode of absorption spectra.

The denaturation curves are well fitted by Eq. 2 (Fig. 5, C-D) but, since BLG HP-denaturation is a pseudo-equilibrium (due to irreversible reactions, see below), the only thermodynamic parameters we extracted are the unfolding volume change (Δ*V_u_*) and the value of the half-denaturation pressure (*P_1/2_*) (Table 2). In agreement with HP-SANS, the UV/vis absorption measurements confirm that retinol strongly stabilizes BLG during HP-denaturation, with a shift in the change of the Trp local structure of ~100 MPa (Fig. 5, B-C) and a significant increased transition pressure (*P*_1/2_) of ~70 MPa in the presence of the ligand (Fig. 5C, Table 2). In contrast, resveratrol has virtually no effect on the stability of the local Trp environment of BLG, as measured by UV/vis absorption (Figs. S5 and 5D, Table 2). The deduced Δ*V_u_* shows that retinol, but not resveratrol, induces a larger volume change during BLG unfolding, as compared to BLG without ligand (Table 2).

**Table 2.** Thermodynamic parameters of pressure denaturation for BLG samples obtained from HP-UV/vis absorption spectroscopy at 30 and 37°C.

| Sample | Temperature (°C) | $\Delta V_u$ (mL/mol) | $P_{1/2}$ (MPa) |
| --- | --- | --- | --- |
| BLG | 30 | $-103 \pm 8$ | $168 \pm 3$ |
| | 37 | $-98 \pm 6$ | $170 \pm 3$ |
| BLG/retinol | 30 | $-158 \pm 9$ | $239 \pm 1$ |
| | 37 | $-191 \pm 15$ | $249 \pm 2$ |
| BLG/resveratrol | 30 | $-114 \pm 8$ | $180 \pm 2$ |

We can therefore conclude that, during *in situ* HP denaturation of BLG, retinol, which has a high affinity binding to BLG cavity, is able to partly but significantly prevent BLG unfolding, both at local Trp environment and at the protein three-dimensional conformation. On the contrary, resveratrol, which binds with low affinity protein surface site(s), has no significant effect on BLG unfolding.

### BLG irreversible refolding and oligomerization

In order to account for BLG concentration effects (5 or 8 mg/mL), SANS data were normalized to protein concentration (Figs. 6A and S6A). After HP treatment up to ~300 MPa, return to atmospheric pressure enables BLG to refold but only partially. At 5 mg/mL, BLG without ligand, while completely unfolded at ~300 MPa (Fig. 3B), is partially refolded at atmospheric pressure, as evidenced by the bell-shaped SANS curves in Kratky representation, characteristic of globular particles (Fig. 6A). As mentioned before, a higher BLG concentration prevents the complete unfolding of the protein, but it may also favor protein partial refolding, as shown by the increased intensity of the peak at 8 compared to 5 mg/ml (Fig. 6A). UV/vis absorption spectroscopy confirms that HP induces irreversible structural local changes, with only ~50% recovery compared to the native protein when pressure returns to atmospheric value (Figs. 5 and S5).

**Figure 6.**
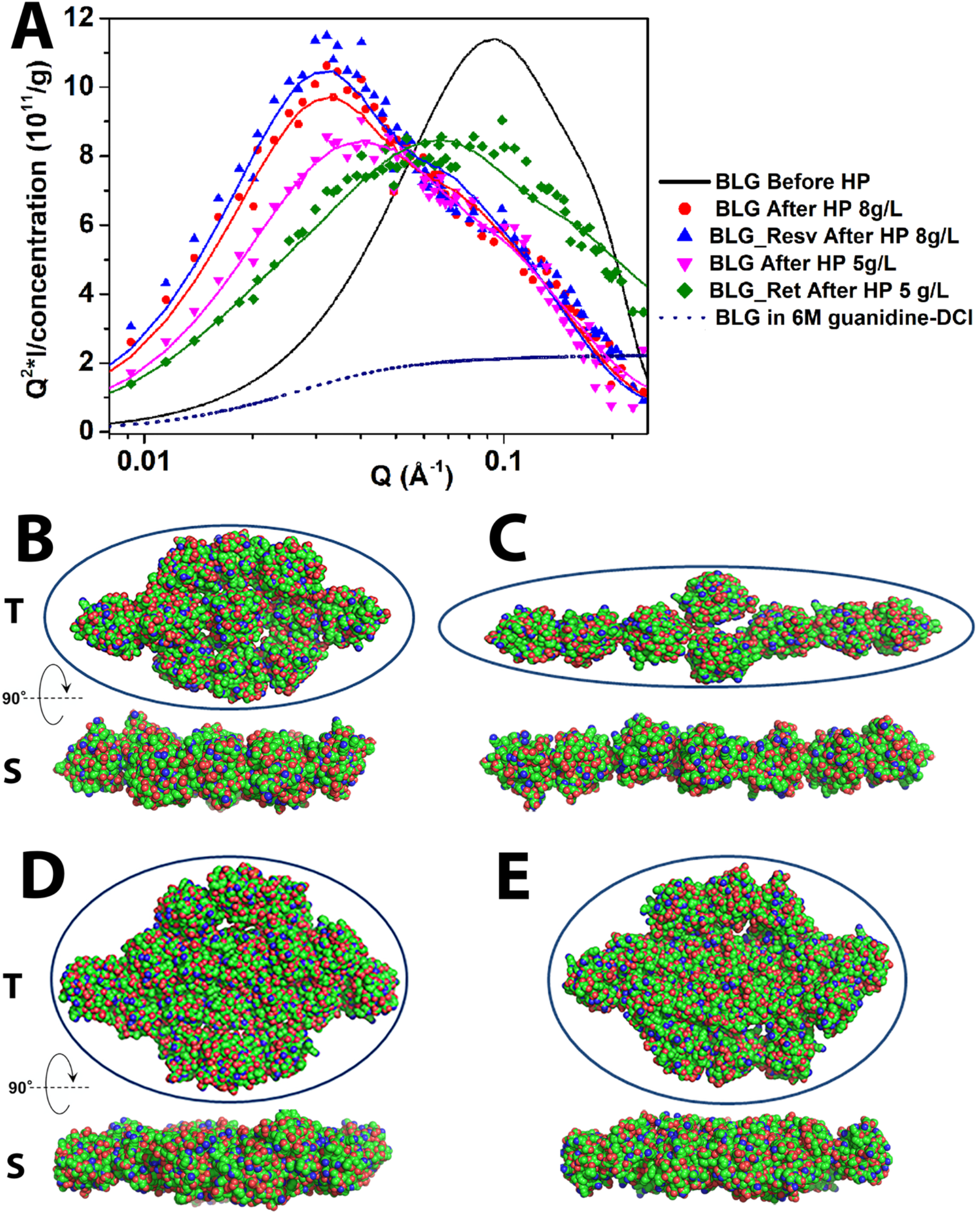
(***A***) Kratky representation of BLG SANS intensities measured, at pD 7.2, in quartz Hellma^®^ cells after ~300 MPa HP treatment and normalized to BLG concentration. Black full and dotted lines are for native or fully chemically unfolded BLG (by deuterated 6 M Gdn-DCl), respectively (8 mg/mL). The other curves represent BLG without ligand at 5 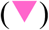 or 8 mg/mL 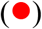, BLG/retinol at 5 mg/mL 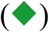, and BLG/resveratrol at 8 mg/mL 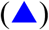. The corresponding colored full lines represent the best fits by the ellipsoid analytical model in SasView program. (***B-E***) *Ab initio* shapes of BLG oligomers corresponding to MASSHA models obtained with the smallest chi^2^ (Table 3). (*B* and *D*) BLG without ligand at 5 or 8 mg/mL, respectively, (*C*) BLG/retinol (5 mg/mL), (*E*) BLG/resveratrol (8 mg/mL). *T* and *S* denote top and side views, respectively. Green, blue, and red spheres represent carbon, nitrogen, and oxygen atoms, respectively. SasView ellipsoid shapes are superimposed to MASSHA model buildings.

Interestingly, HP treatment back and forth up to ~300 MPa causes also irreversible covalent oligomerization, as observed for BLG without ligand at 8 mg/mL, with a large increase of both *I*_*Q*→0_ (from 0.37 to 2.9 cm^−1^) and *R_g_* (from 23.5 to 70.5 Å) (Fig. 6A). We use the term “oligomerization” instead of “aggregation” since the larger objects we observed exhibit a finite size, at least in the SANS *Q*-range we used. At 5 mg/mL, compared to native BLG, the SANS curve after HP treatment is shifted to lower *Q*-values in Kratky representation (from ~0.1 to ~0.04 Å^−1^), which is characteristic of larger scattering objects (Fig. 6A). At 8 mg/mL, the larger size (compared to native protein) of BLG without ligand observed *in situ* at ~300 MPa (Fig. S3) is also increased when pressure returns to 1 bar, as shown by the peak shift from ~0.05-0.06 Å^−1^ (Fig. S3, B & D) to ~0.03 Å^−1^ (Fig. 6A).

The *ab initio* envelopes of BLG oligomers simulated by MASSHA and DAMMIF programs (ATSAS) (30) (34) show that a higher protein concentration induces larger oligomers, which confirms the increased dimensions of the ellipsoid model that fits the data in SasView program (Figs. 6 and S7, B & D, Table 3). MASSHA simulations use BLG crystal structures (monomers or dimers) as building blocks, that are assumed to remain “folded”. However, without making any assumption on BLG folding state, DAMMIF (35) *ab initio* simulations confirm MASSHA results (Fig. S7, B & D).

**Table 3.** Fitting of SANS data using triaxial ellipsoid (SasView) or rigid body modeling (MASSHA, ATSAS). *a*, *b*, *c* represent minor, polar, and major equatorial radii, respectively.

|  | Ellipsoid fitting |  |  |  | Rigid body modeling |  |  |
| --- | --- | --- | --- | --- | --- | --- | --- |
| Sample | $a$ (Å) | $b$ (Å) | $c$ (Å) | $\chi^2$ | Number of<br>BLG<br>monomers | $MW_{app}$<br>(kDa) | $\chi^2$ |
| BLG (5 mg/mL) | 12.8 | 55.0 | 115.5 | 0.73 | 11 | 204.1 | 1.20 |
| BLG/retinol (5 mg/mL) | 8.1 | 35.0 | 163.5 | 0.99 | 8 | 148.4 | 1.45 |
| BLG (8 mg/mL) | 13.4 | 74.4 | 108.9 | 1.68 | 18 | 333.9 | 1.29 |
| BLG/resveratrol (8 mg/mL) | 14.0 | 79.3 | 105.8 | 2.52 | 18 | 333.9 | 1.73 |

### Effect of ligands on BLG oligomerization

Resveratrol binding does not change significantly HP-induced BLG refolding and oligomerization (Fig. 6A). In contrast, retinol binding prevents partly, but significantly, both protein irreversible unfolding and oligomerization extent, as observed with the peak shift to a lower *Q*-value (0.06 Å^−1^) compared to BLG without ligand (0.04 Å^−1^) (Fig. 6A). However, BLG/retinol complex remains significantly different compared to the native dimeric BLG, meaning that HP treatment is not reversible even in the presence of retinol.

MASSHA *ab initio* simulations show that retinol binding reduces the average size of BLG oligomers that are less dense and more elongated compared to BLG without ligand (Fig. 6, B & C, Table 3). DAMMIF simulations confirm that BLG/retinol oligomers exhibit a more linear shape with a smaller density of “beads” (Fig. S7, B & C). In contrast, resveratrol binding does not modify significantly oligomer envelopes obtained by MASSHA (Fig. 6, D & E) and DAMMIF (Fig. S7, D & E).

SDS-PAGE in nonreducing conditions shows that at least one part of HP-induced irreversible oligomerization is covalent in all BLG samples, with the presence of not only covalent dimers but also higher *MW* covalent oligomers (Fig. 7A). A higher BLG concentration induces higher *MW* oligomers in nonreducing SDS-PAGE (Fig. 7B). Resveratrol binding does not significantly decrease the amount of high *MW* oligomers, whereas retinol decreases the amount of both covalent dimers and higher *MW* oligomers, resulting in a higher amount of preserved monomers (Fig. 7).

**Figure 7.**
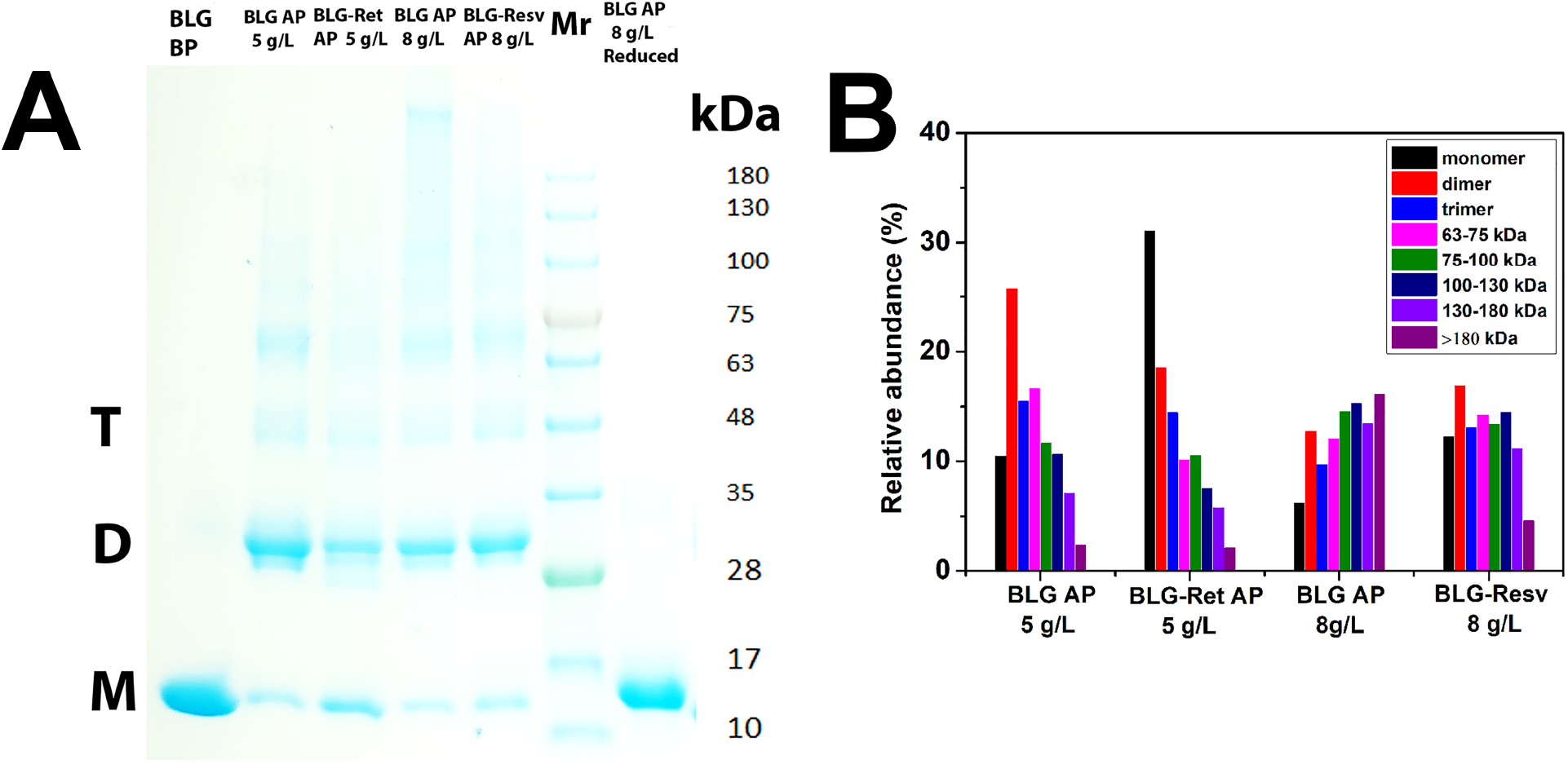
(***A***) SDS-PAGE (4-20% polyacrylamide gel) of BLG samples after HP-treatment at ~300 MPa. (*wells from left to right side*) BLG without ligand before HP, HP-treated BLG without ligand (5 mg/mL), HP-treated BLG/retinol (5 mg/mL), HP-treated BLG (8 mg/mL), HP-treated BLG/resveratrol (8 mg/mL), and HP-treated BLG in reducing conditions (with 5% β-mercaptoethanol) showing the reduction of covalent disulfide bonds. *M, D*, and *T* denote BLG monomers, dimers, and trimers, respectively. *MW* markers (*Mr*) with the corresponding values in kDa are indicated. (***B***) Relative intensities of the protein bands corresponding to the different *MW* of BLG oligomers.

### Temperature effect on BLG structural changes

Since temperature is known to emphasize hydrophobic effects, retinol binding effect on BLG was also measured at 37°C to compare with 30°C (Fig. S8). We found that such a temperature increase strengthens the effect of retinol to prevent HP-induced BLG changes. At ~200 MPa, we observe a reduction of 8.4 Å of the radius of gyration for BLG/retinol complex, compared to that of BLG without ligand, while this reduction is only 4.2 Å at 30°C at the same pressure (Figs. 4B and S9A), as clearly illustrated in Kratky representation (Fig. 3, B & D compared to Fig. S8, C & D). At ~200 MPa, BLG without ligand is much more unfolded by HP at 37°C than at 30°C (Fig. S8E), whereas the effect of retinol on the stabilization of BLG/retinol complex folding is emphasized by temperature (Fig. S8F). *In situ* HP-UV/vis absorption confirms this strengthened effect of retinol on BLG stabilization by temperature. Indeed, *P_1/2_* increases significantly from 239 ± 1 MPa at 30°C to 249 ± 2 MPa at 37°C and the volume reduction is more pronounced, Δ*V_u_* decreasing from −158 ± 9 at 30°C to −191 ± 15 mL/mol at 37°C (Fig. 5C compared to Fig. S10, Table 2). In contrast, a temperature increase from 30 to 37°C has no significant effect on both *P_1/2_* and Δ*V_u_* for BLG without ligand (Table 2).

The unexpected decreased *I*_*Q*→0_ observed upon *in situ* HP denaturation for BLG without ligand at 37°C (Fig. S9B), while at 30°C *I*_*Q*→0_ increases (Fig. S4A), may be due to the limited accessible low *Q*-range in HP cell that prevents to observe large size objects. However, using a larger *Q*-range as in a quartz cell, on return to ambient pressure after ~300 MPa treatment, SANS measurements show that temperature has no or little influence on the aggregated BLG structures (Fig. S6B).

## DISCUSSION

In the present study, we combined *in situ* HP-SANS and HP-UV/vis absorption spectroscopy in an original way to probe the mechanisms of ligand effect on the denaturation of BLG by pressure. We report that retinol, which binds with high affinity BLG cavity, (4) prevents protein unfolding both at 3D and local structural levels and reduces its oligomerization when back to the atmospheric pressure. In contrast, resveratrol, known to bind with a lower affinity on different BLG surface site(s) (7, 33), has virtually no significant effect on BLG structural changes.

Depending on pH, temperature, and concentration conditions, BLG adopts various assembly conformations, *i.e*. monomer, dimer, or octamer structures (14). We checked that, at neutral pH, native BLG is a dimer while the presence of Gdn-DCl, induces its complete unfolding (Figs. 2 and S2), in agreement with structural features observed by SAXS (36).

*In situ* HP-SANS is a powerful tool to probe BLG 3D conformational subtle changes upon pressure increase. No significant change in the structure of BLG is observed up to ~150 MPa, except a slight decrease of *I*_*Q*→0_ probably due to both BLG partial dissociation of native dimers into monomers, as similarly observed by HP-SAXS (11), and an increase of protein hydration, which may reduce BLG contrast in D_2_O buffer. The UV/vis absorption “blue” shift upon pressure, highlighted by spectra 4^th^ derivatives (Figs. 5 and S5), indicates a higher polarity of Trp environment, as a consequence of the movement of the aromatic side chains from the protein hydrophobic environment towards a more hydrated BLG cavity during protein unfolding (9). In particular, Trp19 residue, buried into the hydrophobic cavity of BLG in native condition (Fig. 1), may be exposed to solvent upon denaturation, as previously reported (37).

At 5 mg protein/mL, from ~200 up to ~300 MPa, pressure induces a strong decrease of BLG compactness (Fig. 3, B & D), the continuously reduced SANS intensity being probably due to a higher hydration of the protein, due to both protein unfolding and water penetration inside BLG cavity (38). At that concentration, BLG is completely unfolded at ~300 MPa into a Gaussian chain but its *MW* remains that of the dimer, probably due to disulfide bonds between the chains. The free reactive Cys121 (Fig. 1), not accessible in native BLG, can indeed form intermolecular disulfide bridges when exposed to the solvent by protein unfolding, and also exchange with the disulfide bonds already present in the native state, as reported previously (39).

A higher protein concentration (8 mg/mL) partially preserves BLG from a complete unfolding, as already reported by SAXS and SANS (38), by oligomerization of the native dimers to form tetramers at ~250-300 MPa (Fig. S3). This pair association may not be due to steric (hindrance) effect but rather to disulfide bond formations, as mentioned above. Indeed, at that protein concentration, the average distance between particles (dimers) can be estimated to be ~200 Å, which is shorter than the length of the unfolded protein that is ~150 Å (Fig. 4C).

The high affinity binding of retinol to BLG cavity enables the ligand to significantly stabilize the protein both at tridimensional level (Figs. 3 and 4) and locally at Trp environment (Fig. 5). This is in agreement with a previous study showing that ligand binding to protein cavities increases pressure stability through a higher protein compaction (*40*). A previous HP-fluorescence study showed similar starting pressure points of unfolding for both BLG without ligand and BLG/retinol complex, although this method does not probe directly retinol effects on protein Trp residues but rather retinol fluorescence itself (10). These authors highlight also that retinol dissociates completely from BLG at ~300 MPa (10), in accordance with our SANS data showing at that pressure the total loss of protein compacity (Fig. 3D).

Since BLG denaturation by HP is a pseudo-equilibrium in favor of the unfolded state of the protein, the unfolding volume change Δ*V_u_* we calculated is not the apparent Δ*V^0^* usually deduced from equilibrium reactions. However, Δ*V_u_* is very useful to estimate both conformational transitions and protein/solvent interactions. At the concentration (2 mg/mL) used in the present HP-UV/vis measurements, protein oligomerization is very slow, as observed at 5 mg/mL by *in situ* HP-SANS (Fig. 3). Therefore, we rather focus, by UV-vis absorption spectroscopy, on BLG unfolding and ligand dissociation.

The volume change (−103 mL/mol) we observe for BLG without ligand, at pH 7.2, is roughly comparable to the volume change (−90 mL/mol) at pH 2 obtained by HP-NMR (12). However, this slight difference can be assigned to a reduced resistance of BLG to pressure-induced conformational changes at neutral compared to acidic pH (10).

Apparent Δ*V^0^* is made up of three terms: (*i*) the constitutive volume of atoms, which remains unchanged at the pressures we used, (*ii*) the cavity volume related to protein conformational changes, and (*iii*) the hydration volume taking into account protein/solvent interactions. This third term is often responsible for the negative volume changes induced by HP. For all samples, the calculated Δ*V_u_* are strongly negative, as reported before **(2)**, indicating probably two phenomena. First, a HP-induced increase of the hydration of BLG cavity since the interactions of water molecules with a hydrophobic environment occupy less volume than in bulk water (41–43). The “blue” shift of Trp maximum absorption we observe under HP is compatible with the polarity change of the cavity due to the presence of water binding (Figs. 5 and S5). Second, the electrostriction of water molecules that may interact with the negative charges of BLG residues exposed to the solvent by the protein unfolding (2).

Retinol binding strengthens these observations (relative difference of ~55 mL/mol compared to BLG without ligand, Table 2), suggesting that HP-induced unfolding of BLG presumably dissociates retinol binding from the hydrophobic cavity with the resulting entry of water molecules inside the hydrophobic pocket, like for the protein without ligand. In the case of resveratrol binding, there is a very small negative increase of the volume compared to BLG without ligand (relative difference of ~11 mL/mol, Table 2), since the ligand binding on the BLG surface does not change significantly cavity hydration. Actually, the slight negative increase of Δ*V*_u_, as compared to that of BLG without ligand, may be interpreted as an additional pressure effect on the hydration of the buried surface of BLG/resveratrol complex after dissociation of the ligand (44).

The significant reduction of Δ*V_u_* for BLG/ligand complexes, especially with retinol, upon pressure in comparison to BLG without ligand could also be explained by a larger initial volume of the complex at atmospheric pressure. Volume calculation of BLG crystal structures in the absence or presence of retinol/retinoic acid showed indeed a slightly larger volume (98 mL/mol) of BLG/ligand complexes in comparison to BLG without ligand (45). Similarly, HP-CD study of holo- and apo-myoglobin, a protein with a similar *MW* as BLG, reported a higher volume change with pressure in the presence of ligand (46).

In BLG/*cis*-parinaric acid complex, the ligand, which exhibits a comparable affinity to that of retinol but for an outer hydrophobic binding site (not into the cavity), dissociates at lower pressure than BLG/retinol complex (10). Therefore, ligand stabilization effects on BLG strongly depend on both ligand localization and affinity.

BLG native folding is completely and irreversibly lost by pressures up to ~300 MPa, whether in the absence or presence of ligand. The return to the atmospheric pressure does not enable to recover a complete folded state for BLG but rather a “molten globule” state, as suggested by fluorescence and CD studies showing HP-induced formation of BLG molten globule with long term stability (37). Hence, it seems that an energy barrier between BLG molten globule, stabilized by disulfide bonds, and the native protein cannot be overcome, even in the presence of retinol. HP-induced BLG molten globule exhibits a cavity still able to bind various ligands, but with different affinities in comparison to the native state, whereas HP treatment induces a decrease of retinol binding constant by a factor two, due to conformational changes of the hydrophobic cavity (47). A higher BLG concentration increases apparent *R_g_* and the size of the scattering particles, due to a larger extent of protein oligomerization promoted by HP (Figs. 4 and S3), as already reported by chromatography (48). At 8 mg/mL, BLG structure is “preserved” from a complete unfolding by covalent oligomerization, in agreement with a study showing that thermal aggregation of BLG increases the content of ordered β-sheet secondary structures (49). Our SANS data clearly show that a higher protein concentration contributes to a larger stability of BLG molten globule upon *in situ* HP treatment, probably due to a larger extent of protein covalent aggregation through nonnative disulfide bond exchange, as previously proposed for BLG under HP conditions (37).

HP treatment back and forth up to ~300 MPa produces irreversible oligomerization, as shown by SANS (Figs. 6 and S7) and nonreducing SDS-PAGE (Fig. 7). Similar results were reported by Patel *et al*., at pressures above ~200 MPa using two-dimensional PAGE method (50). Retinol reduces BLG oligomerization probably by a kinetic effect, whereas resveratrol has virtually no effect (Figs. 6 and S7), in agreement with a previous study showing that binding to BLG of high affinity ligands, such as fatty acids, could decrease the size of aggregates induced by heat or HP treatment (51). In retinol-induced stabilization of BLG folding, the protein may spend more time in its native conformation during *in situ* HP treatment, preventing SH group exposure to the solvent and consequently reducing the extent of aggregation. SANS probes both covalent and noncovalent oligomers, giving access to their *MW* (Table 3) and shapes (Figs. 6 and S7) through analytical fitting and *ab initio* simulations. The MASSHA modeling uses the ellipsoid model parameters from SasView (Fig. 6) as constraints to “build” the oligomer shapes, in agreement with the data obtained on BLG oligomerization by gamma irradiation (52). The overall envelopes are also consistent with those obtained from DAMMIF program without any *a priori* geometrical assumption (Fig. S7). Interestingly, we found for BLG/retinol complex the formation of octamers in the presence of retinol (Table 3), as reported in other studies on BLG at pH 4-5 (16). Nonreducing SDS-PAGE gives access to only covalent oligomers, emphasizing the protective effect of retinol and the concentration influence on aggregation, at a qualitative way (Fig. 7A). Unfortunately, the sum of the relative intensities of the protein bands corresponding to the different oligomers (Fig. 7B) is hardly comparable to the apparent *MW* that could be deduced from SANS *I*_*Q*→0_ using Eq. 4, since protein contrast, due to a higher hydration of the protein during unfolding, is unknown.

Although BLG has a relative high thermal stability with *T*_1/2_ of ~70°C (53), HP-induced BLG unfolding is very sensitive to temperature. In particular, the effect of retinol is emphasized by temperature, known to foster hydrophobic interactions (54). For instance, retinol acetate binding to BLG is stimulated by temperature, through rearrangements of the protein secondary structures and side chains interacting with the ligand (55). Our HP-SANS data indicate a higher stabilization effect of retinol at 37°C in comparison to 30°C, as observed for the apparent *R_g_* (Figs. 4B and S9A) and the cylinder form factor parameters (Figs. 5A and S9C), showing that BLG/retinol complex is more compact at 37 than 30°C (Fig. S8F). HP-UV/vis absorption study confirms these results with a higher *P*_1/2_ at 37 compared to 30°C (Figs. 5C and S10). A higher temperature also promotes the hydrophobic aggregation of unfolded BLG induced by HP, producing large oligomers that may not be observed in the restricted *Q*-range of our HP cell, causing an apparent decreased *I*_*Q*→0_ for BLG at 37°C (Fig. S9B) contrary to measurements at 30°C (Fig. S4A).

## CONCLUSION

We report here the effects of two ligands, retinol and resveratrol, with different affinity and binding sites, on the pressure stability of BLG, the main whey protein, by combining *in situ* HP-SANS and HP-UV/vis absorption spectroscopy. Retinol, which binds with high affinity BLG cavity, stabilizes strongly the protein upon HP. Pressure effect on BLG/retinol complex leads to a more negative unfolding volume (Δ*V*_u_), mainly due to the pressure-induced binding of water molecules inside the cavity and ligand dissociation. In contrast, resveratrol, which has a low affinity binding for BLG surface, site(s), shows no significant effect on the stability of the protein, but only a slight negative increase of Δ*V*_u_ compared to BLG without ligand. The return to atmospheric pressure after HP treatment up to ~300 MPa produces irreversible covalent oligomers of BLG, whose shape and size are reduced by retinol, but not resveratrol. More generally, our results highlight the mechanisms involved in the way ligands with high affinity binding for internal hydrophobic cavities stabilize proteins like BLG, especially to resist to pressure denaturation.

## AUTHOR CONTRIBUTIONS

SM, CL, AB, and SC designed the study. SM, BA, AH, CL, AB, and SC conducted and performed HP-SANS experiments. SM, DH, GHBH, and SC conducted HP-UV/vis absorption spectroscopy. SM, AB, and SC analyzed and/or interpreted the data. SM, AB, and SC wrote the article.

## ACKNOWLEDGMENTS

We thank Mikaela Börjesson (LLB, France) for BLG preparation, as well as Adrien Lerbret (PAM, France), Marie-Sousai Appavou (JCNS, Germany), and Raphael Dos Santos Morais (LIBio, France) for their advice. LLB neutron facility is acknowledged for beamtime on PACE SANS instrument.

## Notes

### Competing Interest Statement

The authors have declared no competing interest.

